# Activating neutrophils by co-administration of immunogenic recombinant modified vaccinia virus Ankara and granulocyte colony-stimulating factor for the treatment of malignant peripheral nerve sheath tumor

**DOI:** 10.1101/2023.11.29.569123

**Authors:** Yueqi Wang, Shuaitong Liu, Juan Yan, Shanza Baseer-Tariq, Bharath Salla, Liangliang Ji, Ming Li, Ping Chi, Liang Deng

**Affiliations:** Dermatology Service, Department of Medicine, Memorial Sloan Kettering Cancer Center, New York, NY 10065, USA; Human Oncology and Pathogenesis Program, Memorial Sloan Kettering Cancer Center, New York, NY 10065, USA; Immunology Program, Sloan Kettering Institute, New York, NY 10065, USA; Department of Dermatology, Weill Cornell Medical College, New York, NY 10065

## Abstract

Malignant peripheral nerve sheath tumor (MPNST) is a rare, aggressive soft-tissue sarcoma with a poor prognosis and is insensitive to immune checkpoint blockade (ICB) therapy. Loss-of-function of the histone modifying polycomb repressive complex 2 (PRC2) components, EED or SUZ12, is one of the main mechanisms of malignant transformation. In a murine model of MPNST, PRC2-loss tumors have an “immune desert” phenotype and intratumoral (IT) delivery immunogenic modified vaccinia virus Ankara (MVA) sensitized the PRC2-loss tumors to ICB. Here we show that IT MQ833, a second-generation recombinant modified vaccinia virus Ankara virus, results in neutrophil recruitment and activation and neutrophil-dependent tumor killing in the MPNST model. MQ833 was engineered by deleting three viral immune evasion genes, E5R, E3L, and WR199, and expressing three transgenes, including the two membrane-bound Flt3L and OX40L, and IL-12 with an extracellular matrix anchoring signal. Furthermore, we explored strategies to enhance anti-tumor effects of MQ833 by co-administration of granulocyte colony-stimulating factor (G-CSF).

## Introduction

Malignant peripheral nerve sheath tumor (MPNST) represents an aggressive subtype of soft tissue sarcoma with poor prognosis due to the surgical challenges in local control and lack of effective systemic therapy. MPNSTs occur in distinct clinical settings: type I neurofibromatosis (NF1)-associated (45%), sporadic (45%) or radiation (RT)-associated (10%). They share highly recurrent and biallelic genetic inactivation of three tumor suppressor pathways: *NF1*, *CDKN2A*, and Polycomb repressive complex 2 (PRC2) core components, *EED* or *SUZ12*^1,2,3^. PRC2-loss occurs in more than 80% of all high-grade MPNSTs, and results in global loss of H3K27me2/3 and aberrant transcriptional activation of developmentally silenced master regulators, leading to enhanced cellular plasticity^2^. PRC2-loss in MPNST also leads to aberrant activation of multiple signaling pathways (e.g., WNT signaling), an “immune desert” tumor microenvironment, and primary resistance to immune checkpoint blockade (ICB)^4^.

Viral-based cancer immunotherapy offers a versatile and effective strategy to modulate the immunosuppressive tumor microenvironment (TME) through various mechanisms, such as inducing innate immunity, immunogenic cell death, activation of dendritic cells and T cells, as well as depletion of immunosuppressive cells^5–9^. The TME consists of diverse myeloid cells and lymphocytes, whose antitumor functions are often impaired by the hostile environment of growing tumors^10–12^. Reprogramming myeloid cells and reinvigorating lymphocytes to exert potent and persistent antitumor activities are major challenges in cancer immunotherapy^13–15,16,17,18^. Immunogenic viruses have emerged as a strategy to alter the TME and enhance antitumor immunity^19–23,24–26^.

Poxviruses are a family of large cytoplasmic DNA viruses. Among them, modified vaccinia virus Ankara (MVA) stands out as a highly attenuated vaccinia virus extensively used as a vaccine vector^24–26^. MVA infection of dendritic cells induces type I IFN through the cytosolic DNA-sensing pathway, mediated by cyclic GMP-AMP synthase (cGAS) and downstream signaling molecules such as Stimulator of Interferon Genes (STING)^27^. However, MVA encodes multiple inhibitors of the nucleic acid-sensing pathways. Intratumoral (IT) delivery of heat-inactivated MVA (Heat-iMVA) generates stronger antitumor immunity than live MVA. This response relies on requires CD8^+^ T cells, Batf3-dependent CD103^+^/CD8a cross-presenting dendritic cells (DCs), and STING-mediated cytosolic DNA-sensing pathway^22^.

To improve MVA-based cancer immunotherapy, we have now engineered first and second generations of recombinant MVA viruses. Our first-generation recombinant MVA (rMVA/MQ710) was engineered to delete a viral immune evasion gene E5R, which encodes a potent inhibitor of cGAS^28^, and to express two transgenes including Flt3L and OX40L^29^. IT rMVA/MQ710 (MVAΔE5R-Flt3L-OX40L) generates potent antitumor immunity, dependent on CD8^+^ T cells, the cGAS/STING, and type I IFN signaling. Remarkably, IT rMVA/MQ710 depletes OX40^hi^ regulatory T cells via OX40L/OX40 interaction and IFNAR signaling^29^.

To enhance viral-mediated antitumor effects, we engineered our second-generation recombinant MVA (MQ833) with the deletion of two more viral immune evasion genes – E3L and WR199, and the insertion of IL12 anchored to the extracellular matrix to mitigate toxicity^30^. IT MQ833 demonstrated remarkable antitumor efficacy, resulting in high cure rates in murine tumor models including B16-F10 and MC38. MQ833 also showed efficacy in ICB-resistant murine melanomas with MHC-I loss^30^.

In this study, we explored the use of MQ833 for the treatment of “immune-desert” PRC2-loss MPNST and found that PRC2-loss tumors had better response to MQ833 than PRC2-wt tumors. We also observed that neutrophils, macrophages, CD4^+^, and CD8^+^ T cells play critical roles for virus-induced antitumor effects. Single-cell RNA sequencing (scRNA-seq) of CD45^+^ tumor-infiltrating immune cells in MPNST tumors treated with MQ833 revealed recruitment and activation of neutrophils and monocytes and reprogramming tumor-infiltrating myeloid cells and T cells. Furthermore, co-administration of MQ833 and human G-CSF enhanced antitumor effects. Taken together, these results provide strong preclinical data and rationale for the use of the combination strategy for the treatment of MPNST in the clinic.

## Results

### MQ833 infection promotes type I IFN production in murine MPNST cell lines

Our previous results demonstrated that IT delivery of heat-inactivated modified vaccinia virus Ankara (MVA) enhanced tumor immune infiltrates and sensitized PRC2-loss tumors to ICB^1^. To improve therapeutic efficacy, we have designed our first and second generations recombinant MVA viruses (MQ710 and MQ833) by expressing transgenes to modulate TME and by inducing type I IFN from immune and tumor cells. We found that IT MQ833 generated stronger antitumor effects than MQ710 in murine B16-F10 melanoma and MC38 colon cancer model^30^. We focused on MQ833 for the MPNST model using a well-characterized murine MPNST tumor-derived cell line SKP605 (*Nf1^-/-^Cdkn2a^-/-^ Cdkn2b^-/-^*) from skin-derived precursors (SKPs) of the C57BL/6J mice^31,32^. Two isogenic cell lines, SKP605 sg*Con* (PRC2-wt) and SKP605 sg*Eed* (PRC2-loss) were generated by using CRISPR-Cas9 to knock out the PRC2 core component *Eed*^4^. As previously described^30^, MQ833 was generated by deleting of both E3L and WR199 genes from the MQ710 genome and inserting of murine IL-12 expression cassette in the WR199 locus (Fig. 1a). Vaccinia E3L gene encodes a 190 amino acid protein (E3) with an N-terminal Z-NA (nucleic acid) binding domain and a C-terminal dsRNA-binding domain^33^. E3 antagonizes dsRNA-sensing pathway mediated by MDA5/MAVS^34^. WR199 encodes a 68-KDa ankyrin repeat/F-box protein^35^, another cGAS inhibitor^28^.

**Fig. 1.**
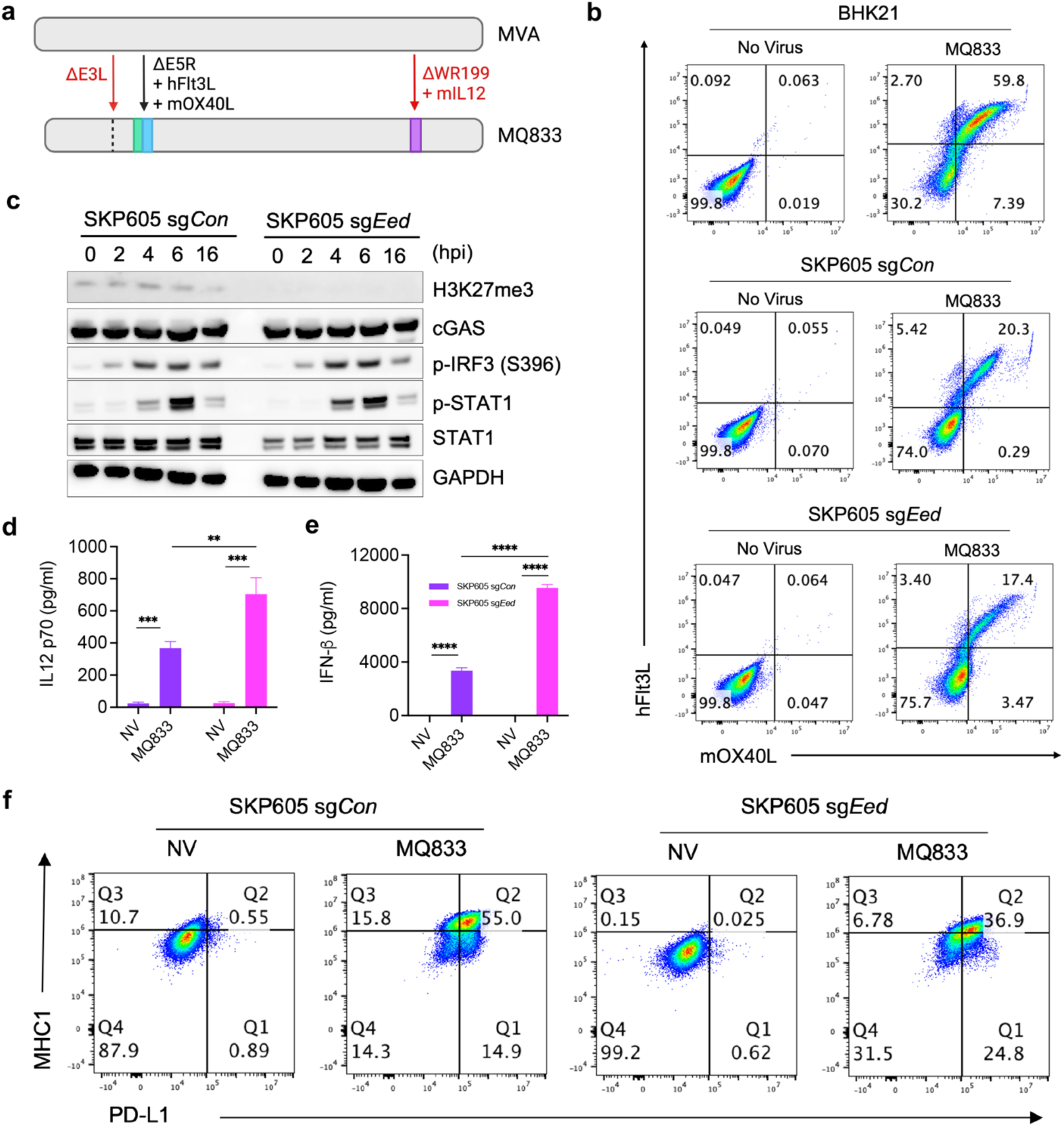
MQ833 infection promotes type I IFN production in murine MPNST cell lines. **a,** Diagram of MQ833 design. (**b-e)**, SKP605 sg*Con* and sg*Eed* cells were infected by MQ833 at a MOI of 10. Cells and supernatant were collected at 16 hours post infection (hpi). **b**, Flow cytometric analysis of mOX40L and hFlt3L transgene expression. SKP605 sg*Con* and sg*Eed* cells were infected with MQ833 at a MOI of 10, and cells were collected at 2, 4, 6, 8, 16 hours post infection (hpi). **c,** Western blot for indicated protein and phosphorylated proteins. **d**, ELISA test for mIL-12 p70. **e**, ELISA test for mIFN-β. **f**, Flow cytometric analysis of MHC1 and PD-L1 expression. SKP605 sg*Con* and sg*Eed* cells were infected with MQ833 at a MOI of 10, and cells were collected after infection for 16 hours.

We first charactered the infection of MQ833 in SKP605 sg*Con* (PRC2-wt) and SKP605 sg*Eed* (PRC2-loss) cell lines *in vitro*. Flow cytometry analyses showed the expression of mOX40L and hFlt3L on the surface MQ833-infected SKP605 sg*Con* and sg*Eed* cells (Fig. 1b). Western blot analyses confirmed the lack of H3K27me3 in SKP605 sg*Eed* cells (Fig. 1c) and showed that MQ833 infection induced the phosphorylation of IRF3 and STAT1 in both cell lines (Fig. 1c). In addition, MQ833 infection triggered the secretion of both IL-12 and IFN-β, with higher levels of both cytokines in the supernatants from SKP605 sg*Eed* cells compared to those from SKP605 sg*Con* cells (Fig. 1d, e). These results indicate that MQ833 potently induces IFN-β secretion from SKP605 cells, which is likely due to the activation of both the cytosolic DNA and dsRNA-sensing pathways^30^. In addition, we observed that MQ833 infection induces MHC-I and PD-L1 expression in both SKP605 sg*Con* and sg*Eed* cell lines (Fig. 1f).

### IT MQ833 generates potent antitumor effects in PRC2-wt and PRC2-loss murine MPNST model, synergizing with ICB therapy using anti-CTLA-4 and anti-PD-1 antibodies

We then evaluated the antitumor effects of MQ833 in murine MPNST models, with or without co-administration of ICB inhibitors. Briefly, SKP605 sg*Con* or sg*Eed* cells were implanted subcutaneously into the right flanks of C57BL/6J mice. Once the tumors were established, they were treated with either IT MQ833, IT MQ833 plus i.p. ICB (intraperitoneal delivery of anti-CTLA-4 and anti-PD-1 antibodies), or with PBS twice weekly. Tumor sizes were measured, and mouse survival was monitored (Fig. 2a). IT MQ833 generated stronger antitumor effects in PRC2-loss SKP605 sg*Eed* tumors compared with PRC2-wt SKP605 sg*Con* tumors (62.5% vs. 0%; *P* < 0.001). The combination of IT MQ833 and systemic delivery of anti-CTLA-4 and anti-PD-1 antibody resulted in 100% cure in the PRC2-loss SKP605 tumor and 75% cure in the PRC2-wt tumor (Fig. 2b, c). These results demonstrate that IT delivery of MQ833 generates potent antitumor effects in PRC2-wt and PRC2-loss MPNST tumors, overcoming ICB resistance, and that combination therapy with ICB inhibitors further enhances the anti-tumor effects.

**Fig. 2.**
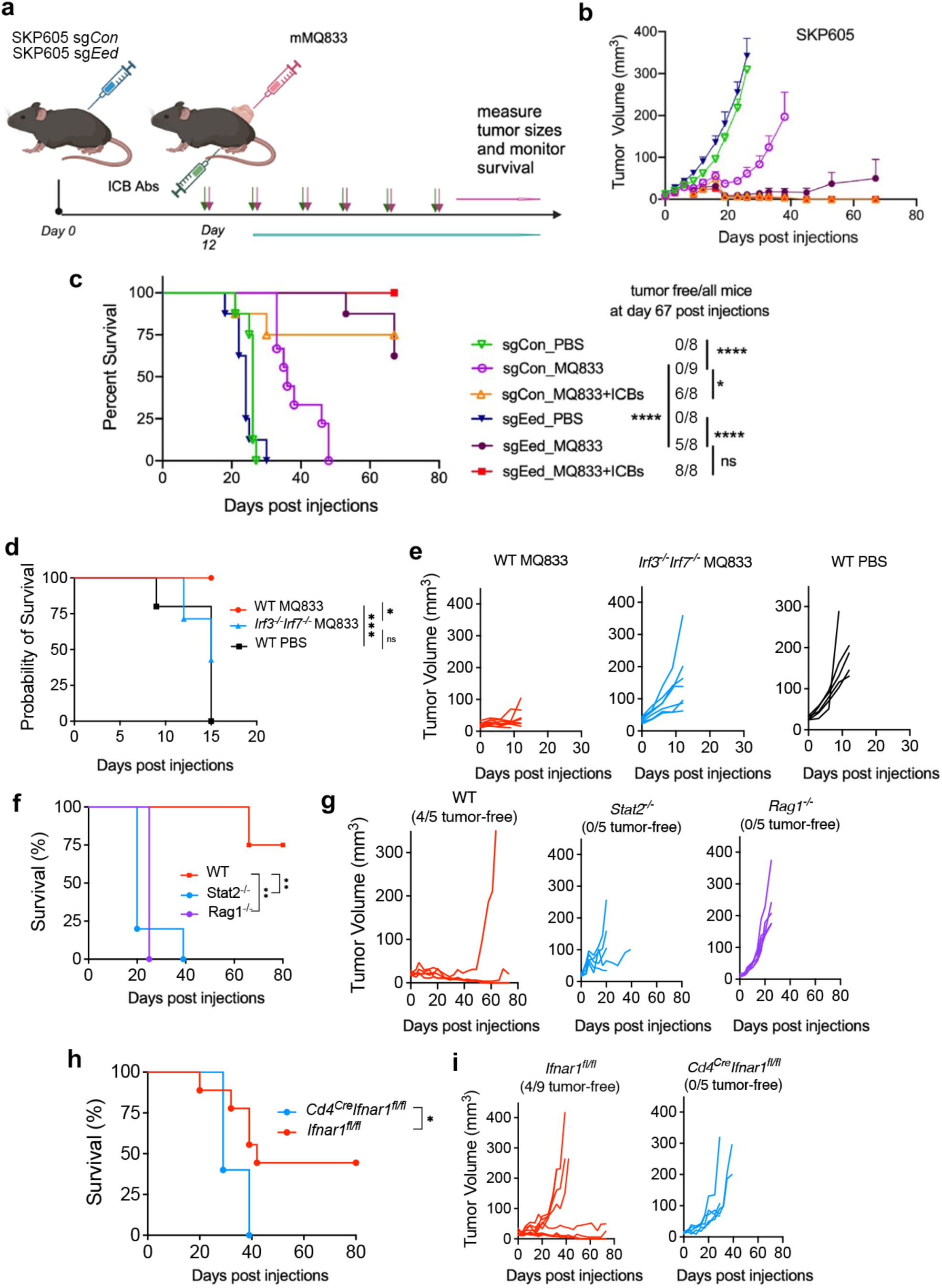
IT MQ833 generates potent antitumor effects in MPNST model, synergizing with ICB therapy. **a,** Schematic diagram of the experimental schedule. 1×10^6^ of SKP605 sg*Con* or SKP605 sg*Eed* cells were subcutaneously implanted into the right flank of C57BL/6J mice. Intratumoral virus treatments were given twice a week once the tumors were established. Tumor size and mice survival were monitored. **b,** Tumor volumes of mice injected with MQ833 or PBS over days post injection (DPI). **c,** Kaplan-Meier survival curve of MPNST tumor bearing mice treated with MQ833 virus. PBS was used as a control. (n=8-9) Survival data were analyzed by log-rank (Mantel-Cox) test. *, P < 0.1. **(d-i)** 1×10^6^ of SKP605 sg*Eed* cells were subcutaneously implanted into the right flank of *Irf3^-/-^ Irf7^-/-^, Stat2^-/-^, Rag1^-/-^, Cd4^cre^Ifnar1^fl/fl^* and WT C57BL/6J mice. Intratumoral virus treatments were given twice a week once the tumors were established. Tumor size and mice survival were monitored. Kaplan-Meier survival curve and tumor volume of MPNST tumor bearing Irf3^-/-^ Irf7^-/-^ mice (**d,e**), Stat2^-/-^ mice (**f, g**), Rag1^-/-^ mice (**f, g**), and *Cd4^cre^Ifnar1^fl/fl^* mice (**h, i**), treated with MQ833 virus. PBS and WT mice was used as a control. Survival data were analyzed by log-rank (Mantel-Cox) test. *, P < 0.1.

### IT MQ833-induced antitumor effects are dependent on type I IFN production, IFNAR signaling and adaptive immunity in a murine MPNST model

We have previously shown that MQ833 infection of murine bone marrow-derived dendritic cells (BMDCs) induced high levels of type I IFN production and the anti-tumor effects were reduced in mice deficient of *Mda5* and *Sting*, adaptors of the cytosolic dsRNA- and DNA-sensing pathways, or in mice deficient of *Stat2*, indicating that type I IFN production and IFNAR signaling mediated by *Stat2* are critical for MQ833-induced antitumor effects^30^.

To investigate whether similar mechanism holds true for the MPNST tumor model, we implanted SKP605 sg*Eed* cells into the right flanks of WT, *Irf3*^-/-^*Irf7*^-/-^, and *Stat2*^-/-^ mice, and found that IT MQ833 resulted in 80-100% cure in WT mice but no cure in *Irf3*^-/-^*Irf7*^-/-^ or *Stat2*^-/-^ mice (Fig. 2d, e, f). IRF3 and IRF7 are two critical transcription factors regulating type I IFN expression. STAT2 is an essential transcription factor downstream of IFNAR signaling. These results confirmed the importance of IFN production and IFNAR signaling in MQ833-mediated anti-tumor effects in a murine MPNST model.

To test whether the adaptive immune system is essential for MQ833-induced antitumor immunity, we used *Rag1*^-/-^ mice, which lack mature B and T cells. IT MQ833 in *Rag1*^-/-^ mice failed to control tumor growth and all mice died, with a median survival of 25 days (Fig. 2f, g).

To test whether IFNAR1 signaling on T cells plays a crucial role for MQ833-induced antitumor immune responses, we generated *Cd4*^cre^*Ifnar1*^fl/fl^ mice in which *Ifnar1* is conditionally knocked-out only in CD4 and CD8 T cells. IT MQ833 eradicated tumors in 4 out of 9 *Ifnar1*^fl/fl^ mice, however, none of the *Cd4*^cre^*Ifnar1*^fl/fl^ mice were cured with the same treatment (Fig. 2h, i).

IFNAR signaling was shown to be important in MQ833-induced antitumor effects in a B16-F10 melanoma model^30^. To test whether IFNAR1 signaling on neutrophils is important, we generated *Mrp8*^cre^*Ifnar1*^fl/fl^ mice in which *Ifnar1* is conditionally knocked-out only in neutrophils. Our results showed larger initial tumor sizes in *Mrp8*^cre^*Ifnar1*^fl/fl^ mice than those in the *Ifnar1*^fl/fl^ mice (Supplementary Fig. 1a), failed to respond to MQ833 treatment and died around 10 days post tumor implantation (Supplementary Fig.1b, c). Taken together, these results indicate that IFNAR1 signaling on T cells and neutrophil are critical for MQ833-induced antitumor effects.

### The antitumor effects of MQ833 are dependent on neutrophils, macrophages, CD4^+^, and CD8^+^ T cells

We performed antibody depletion experiments to determine which immune cells were involved in the MQ833-induced antitumor effects in the PRC2-loss MPNST model (Fig. 3a). SKP605 sg*Eed* cells (1 x 10^6^) were implanted subcutaneously into the right flanks of C57BL/6J mice. Once the tumors are established, immune cell subsets were depleted by administering 200 μg monoclonal antibodies i.p. twice weekly starting 1 day prior to the first viral injection as indicated, continued until the animals either died, were euthanized, or were completely clear of tumors: anti-CD8-α for CD8+ T cells (clone 2.43, BioXCell), anti-CD4 for CD4^+^ T cells (clone GK1.5, BioXCell), anti-NK1.1 for NK cells (clone PK136, BioXCell), and 500 μg of anti-Ly6G (clone 1A8, BioXCell) for neutrophil depletion. 500 μg of anti-CSF1 (provided by Ming Li’s laboratory) were given every 5 days for macrophage depletion. Tumor sizes were measured, and mice survival were monitored (Fig. 3a). Depleting neutrophils via treatment with anti-Ly6G antibody significantly decreased antitumor efficacy, whereas depleting macrophages via anti-CSF1 antibody treatment had a modest impact (Fig. 3b, c). Surprisingly, depleting CD4 cells had a detrimental effect on treatment efficacy, whereas CD8 depletion caused a modest reduction of antitumor efficacy (Fig. 3b, c). However, depletion of NK cells slightly improved MQ833-induced antitumor effects (Fig. 3b, c). These results demonstrated that IT MQ833 activates both the innate and adaptive immunity, including neutrophils, macrophages, CD4^+^ and CD8^+^ T cells to facilitate tumor killing in a PRC2-loss MPNST model, which is resistant to ICB therapy.

**Fig. 3.**
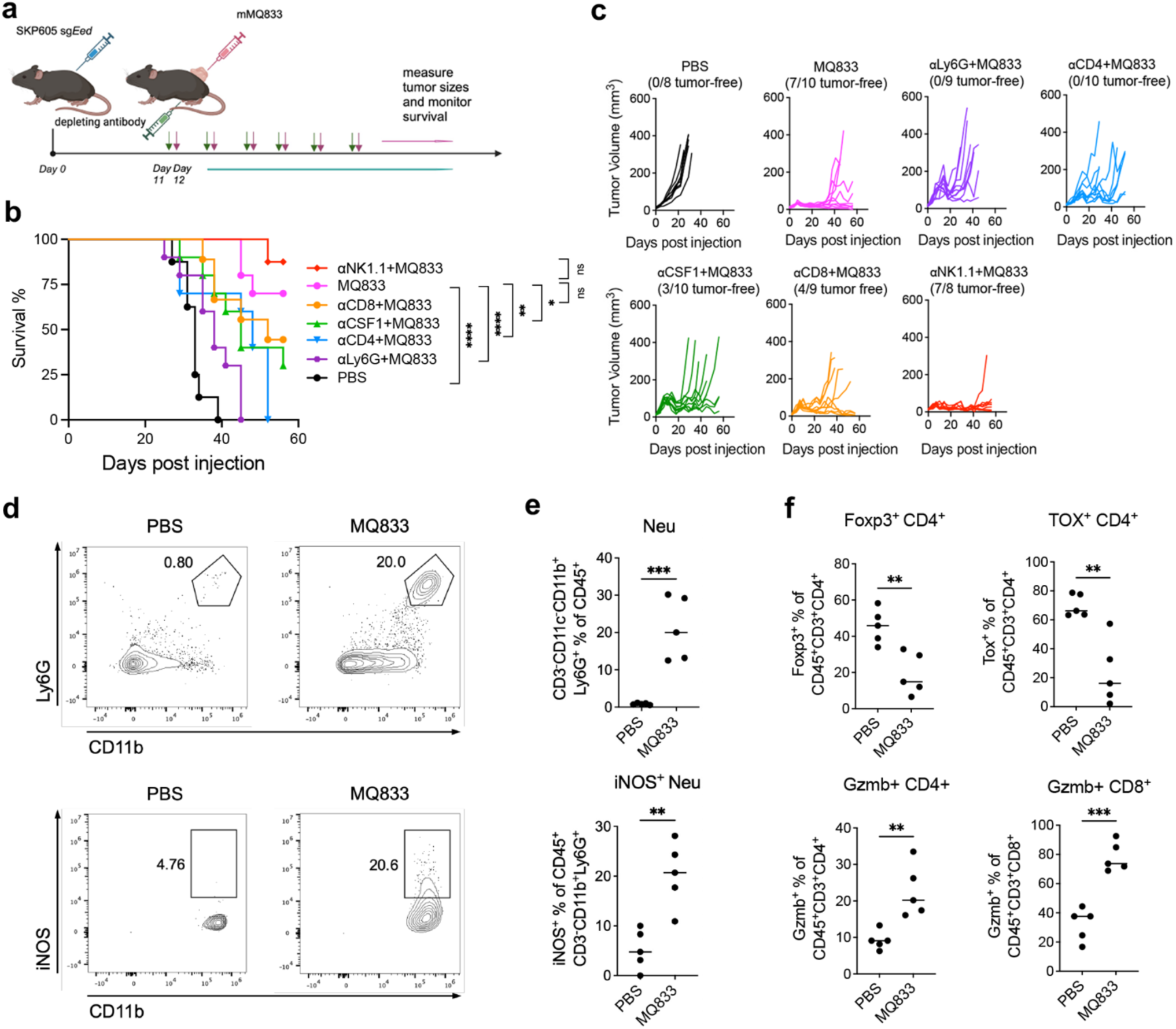
The recruitment and activation of CD4 T cells and neutrophils are crucial for the antitumor efficacy induced by MQ833. **a,** Schematic diagram of the experimental schedule. 1×10^6^ of SKP605 sg*Eed* cells were subcutaneously implanted into the right flank of WT C57BL/6J mice. IT MQ833 injections were given twice a week once the tumors were established. I.p. αCD8, αCD4, αCSF1, αNK1.1, and αLy6g antibodies were given one day prior to every virus injection. I.p. αCSF1 were given every 7 days starting at one day prior to the first virus injection. Tumor size and mice survival were monitored. **b,** Kaplan-Meier survival curve of mice in each treatment group. PBS was used as a control. (n=8-10) Survival data were analyzed by log-rank (Mantel-Cox) test. *, P < 0.05. **c**, Tumor volumes of mice separated by group over days post treatment. Contour plots (**d**) and bar graphs (**e**) showing percentage of CD45^+^CD3^-^ CD11b^+^Ly6g^+^ neutrophils among CD45^+^ Cells (top row), and the percentage of iNOS^+^ cells among CD45^+^CD3^-^CD11b^+^Ly6g^+^ neutrophils (bottom row) in MQ833 or PBS treated tumors. **f,** Percentage of FoxP3^+^, TOX^+^, Gzmb^+^ CD4^+^ or CD8^+^ cell population in MQ833 or PBS treated tumor.

### IT MQ833 activates neutrophils, reverses T cell exhaustion, and depletes Tregs in MPNST tumors

We investigated the ability of IT MQ833 treatment to alter the immunosuppressive tumor microenvironment (TIME) of PRC2-loss MPNST tumors. SKP605 sg*Eed* (1 x 10^6^) cells were implanted subcutaneously into the right flanks of C57BL/6J mice. On day 12 post implantation, when the tumors were established, IT MQ833 (8 x 10^7^ pfu) was injected into the tumors twice, three days apart. Tumors were harvested and single cells were prepared by enzymatic digestion and stained for immune cell analyses. IT MQ833 resulted in neutrophil recruitment and activation with iNOS upregulation in PRC2-loss MPNST tumors (Fig. 3d, e). The percentages of neutrophils (Ly6G^+^CD11b^+^CD45^+^) out of CD45^+^ cells in the IT MQ833-treated tumors increased from 1% to about 20% (Fig. 3d, e). In addition, the percentage of iNOS^+^ neutrophils increased from 5% to 21% (Fig. 3d, e).

Flow cytometry analyses of the tumor-infiltrating T cells revealed that IT MQ833 in PRC2-loss SKP605 tumors resulted in increased expression of the activation marker Granzyme B (Fig. 3f) and decreased expression of the exhaustion marker Tox in CD8^+^ T cells (Fig. 3f). Similarly, upregulation of Granzyme B expression (Fig. 3f) and down-regulation of Tox expression was also observed in CD4^+^ T cells (Fig. 3f). In addition, IT MQ833 resulted in dramatic reduction of Foxp3^+^ CD4^+^ T cells in the treated tumors (Fig. 3f).

Taken together, these results demonstrated that IT MQ833 triggered neutrophil recruitment and activation, as well as CD4^+^ and CD8^+^ T cell activation and the depletion of Tregs.

### Single cell RNA sequencing (scRNA-seq) of tumor-infiltrating immune cells before and after IT MQ833 treatment reveals an alteration of TME and a broad induction of type I IFN pathway

To gain a comprehensive and unbiased understanding of the transcriptome profiles of immune cell populations in SKP605 sg*Con* and sg*Eed* tumors prior to and post IT MQ833, we isolated tumors two days after the second injection of MQ833 and sorted CD45^+^ cells from single cell suspensions and performed scRNA-seq. Unsupervised clustering revealed a total of 21 distinct clusters among the tumor-infiltrating CD45^+^ immune cells (Supplementary Fig. 2)., including NK (C0), Tcf7^+^ T cells (C6), memory T cells (C11), proliferating T cells (C15 and C9), CD4^+^ T cells including Tregs (C3), M1-like macrophages/monocytes (C4 and C8), M2-like macrophages (C1, C2, and C7), neutrophils (C5 and C19), B cells (C14 and C18), conventional DC2 (cDC2) (C10), migratory DCs (C13), plasmacytoid DCs (C16), and basophils (C17) (Fig. 4a-d). IT MQ833 resulted in profound changes in the immune cell compositions, including recruitment of neutrophils (C5 and C19), expansion of monocytes/M1-like macrophages (C4 and C8), reduction of M2-like macrophages (C1 and C2), reduction of Tregs (C3), and expansion of proliferating T cells (C15 and C9) (Fig. 4a-d). We also observed a broad upregulation of interferon stimulated genes (*Isg15*) in myeloid cells after virus treatment (Fig. 4c).

**Fig. 4.**
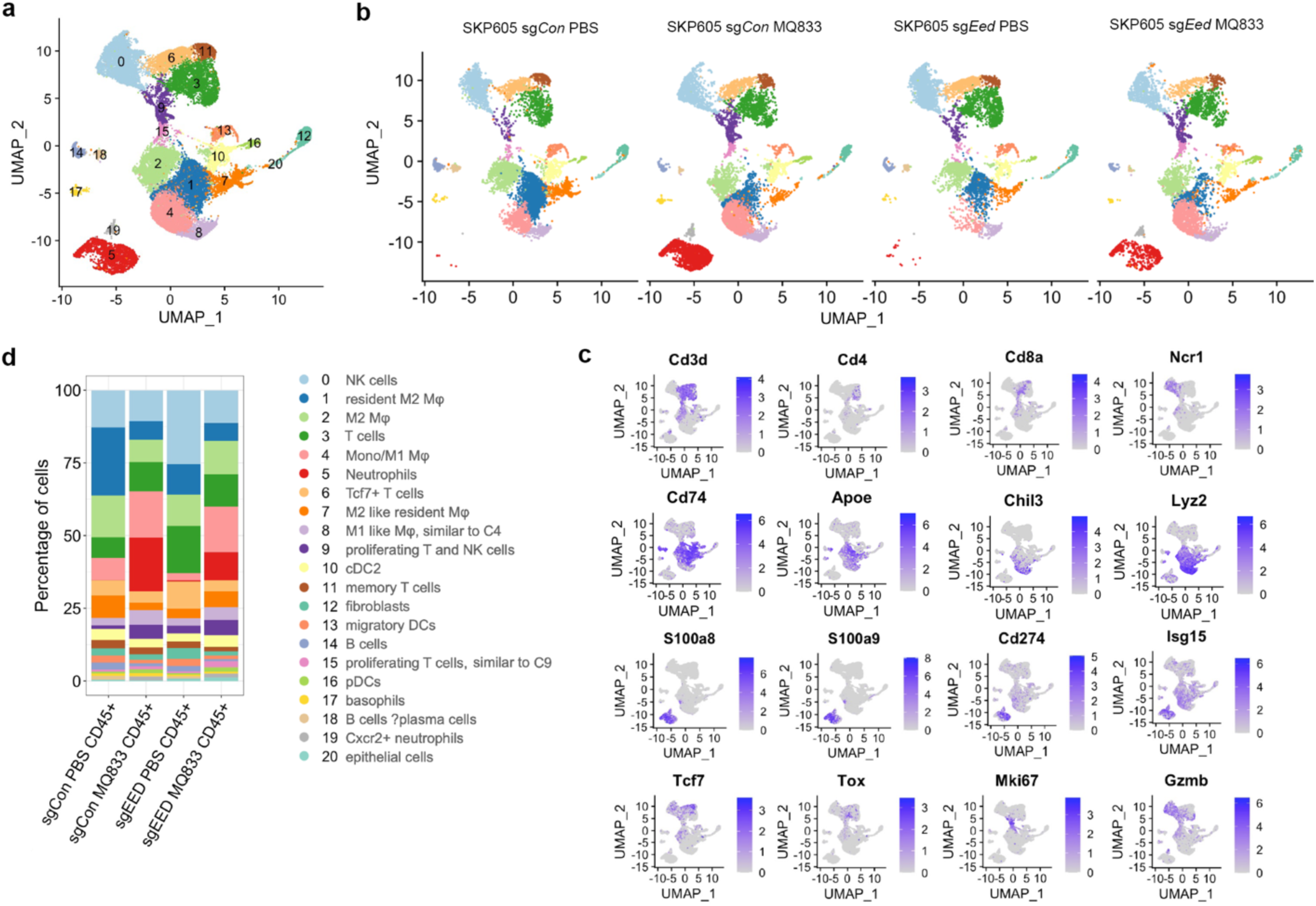
IT MQ833 injection altered TME and broadly induced type I IFN pathway. 1×10^6^ of SKP605 sg*Con* or SKP605 sg*Eed* cells were subcutaneously implanted into the right flank of C57BL/6J mice. One IT MQ833 injection was given once the tumors were established. Tumors were collected and processed into single cell suspension, and CD45^+^ cells were sorted and scRNA-seq analysis were performed. **a,** UMAP display of sorted CD45^+^ cells from 4 samples combined following 10X Genomics scRNA-seq workflow (n = 33201 cells). **b,** UMAP plots of cell clusters separated by sample. **c,** Expression plots of main cell types using their top marker genes. **d,** Stacked bar plot showing the percentage of each cluster across different samples.

### Sub-clustering analyses of tumor-infiltrating T cells reveals IT MQ833 depletes Tregs and promotes the expansion of ISG^+^ CD4+, effector CD8^+^, and proliferating CD4^+^ and CD8^+^ T cells

Sub-clustering analysis of CD3^+^ T cells revealed 11 distinct T-cell clusters (Supplementary Fig. 3), including NK-like T Cells (C0), ISG^+^ CD4^+^ T cells (C1), Tregs (C2), Stem-like T cells (C3), Effector CD8^+^ T cells (C4), Pdcd1^+^Lag3^+^CD8^+^ T Cells (C5), proliferating T cells (C6, C10), Lyz^+^CD3^+^ T cells (C7), ψο T cells (C9) (Fig. 5a, b). Notably, IT MQ833 treatment induced a decrease in Tregs and an increase in ISG^+^CD4^+^ T cells, effector CD8^+^ T cells, and proliferating T cells. PRC2-loss tumors have a higher percentage of exhausted Pdcd1^+^Lag3^+^CD8^+^ T Cells than PRC-wt tumors (C5) (Fig. 5c). Remarkably, there was a notable reversal of the exhaustion following MQ833 treatment in PRC2-loss tumors (Fig. 5c). These results provide transcriptomic details illustrating the impact of MQ833 treatment on remodeling of the T cell compartments in the treated tumors.

**Fig. 5.**
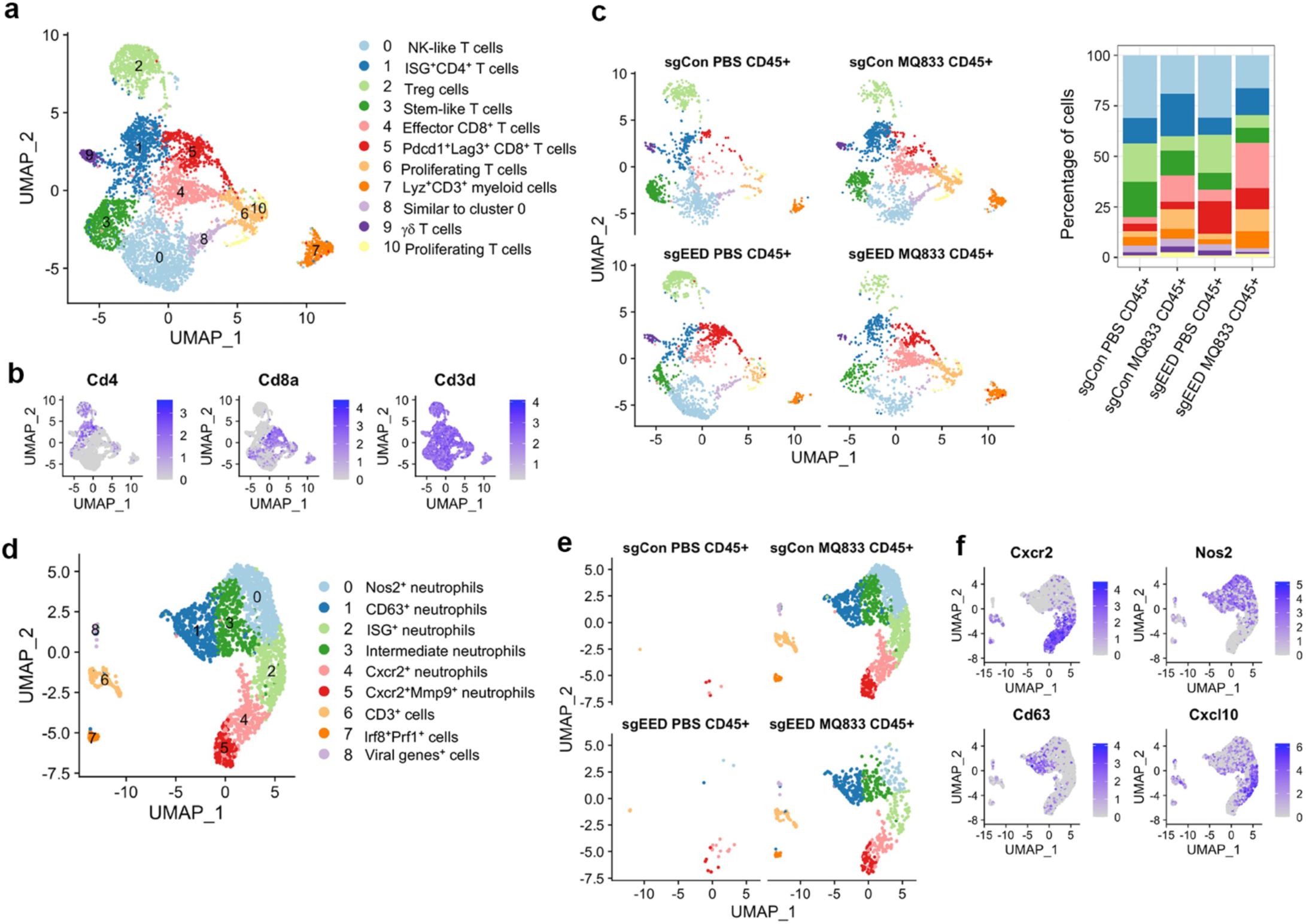
IT MQ833 delivery remodels the T cells and neutrophils of TME. 1×10^6^ of SKP605 sg*Con* or SKP605 sg*Eed* cells were subcutaneously implanted into the right flank of C57BL/6J mice. One IT MQ833 injection was given once the tumors were established. Tumors were collected and processed into single cell suspension, and CD45^+^ cells were sorted and scRNA-seq analysis was performed. **a,** UMAP display of subclustered CD45^+^CD3^+^ cells from 4 samples combined following 10X Genomics scRNA-seq workflow (n = 5,943 cells). **b,** Expression plots of *Cd4, Cd8a, Cd3d.* **c,** UMAP plots and stacked bar plot of cell clusters separated by sample. **d,** UMAP display of subclustered neutrophils from 4 samples combined following 10X Genomics scRNA-seq workflow (n = 2,399 cells). **e,** UMAP plots of cell clusters separated by sample. **f,** Expression plots of *Cxcr2, Nos2, Cd63, Cxcl10*.

### Sub-clustering analyses of tumor-infiltrating neutrophils reveals subsets of neutrophils with distinctive markers upon MQ833 treatment

IT MQ833 in both PRC2-wt and PRC2-loss MPNST resulted in a large influx of neutrophils (Fig. 4) compared with PBS-treated tumors. A sub-clustering analysis of tumor-infiltrating neutrophils revealed six sub-populations (Supplementary Fig. 4)., including Nos2^+^ neutrophils (C0), CD63^+^ neutrophils (C1), ISG^+^ neutrophils (C2), intermediate neutrophils (C3), Cxcr2^+^ neutrophils (C4), and Cxcr2^+^Mmp9^+^ neutrophils (C5). Notably, the activated neutrophil subclusters, namely Nos2^+^ (C0), CD63^+^ (C1), and the intermediate (C3), displayed scarce expression of Cxcr2, while the Cxcr2^+^ (C4) and (C5) lacked Nos2 expression (Fig. 6f). ISG^+^ neutrophils (C2) had the highest level of Cxcl10 expression compared with the rest of the subsets (Fig. 6f). These results demonstrate the heterogeneity of neutrophils in the TME after MQ833 treatment. Nos2 encodes an inducible nitric oxide synthase (iNOS), responsible for the production of nitric oxide (NO), which has pleotropic effects including context-dependent protumor and antitumor functions and antimicrobial activities^36,37,38^. The generation of Nos2^+^ neutrophils are likely induced by IFNGR1 signaling on neutrophils^30^.

**Fig. 6.**
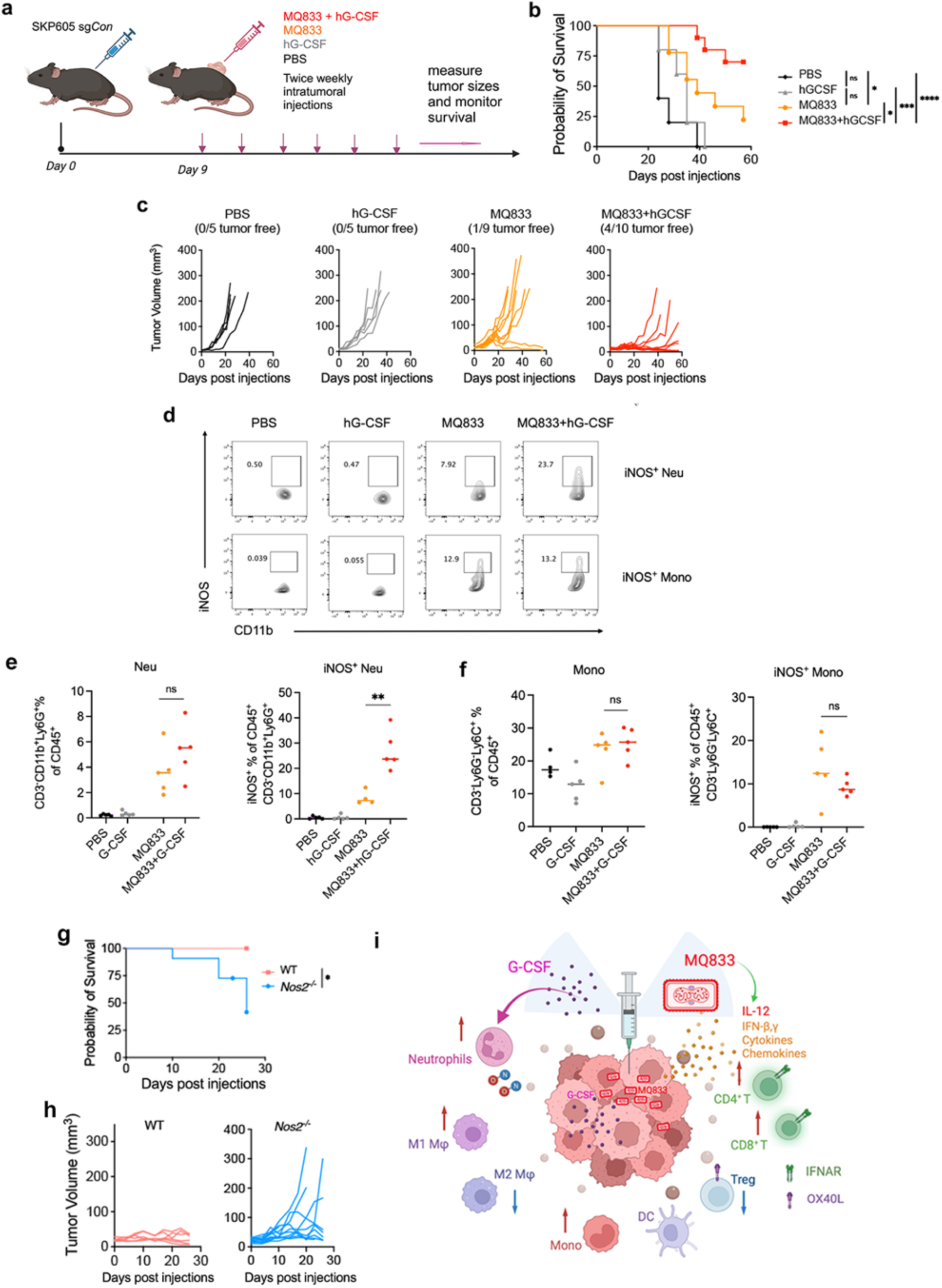
Co-injection of MQ833 and human G-CSF enhances the therapeutic efficacy in MPNST by promoting iNOS expression in tumor-infiltrating neutrophils. **a**, 1×10^6^ of SKP605 sg*Con* cells were subcutaneously implanted into the right flank of WT C57BL/6J mice, IT MQ833 injections, with or without 3µg G-CSF, were given twice a week once the tumors were established. Tumor size and mice survival were monitored. **b,** Kaplan-Meier survival curve of SKP605 sg*Con* tumor bearing mice treated with MQ833 alone or with G-CSF. PBS was used as a control. (n=5-10) Survival data were analyzed by log-rank (Mantel-Cox) test. *, P < 0.1. **c,** Tumor volumes of mice with indicated treatment over days post treatment. **d**, Contour plots (**e,f**) and bar graphs showing percentage of iNOS^+^ cells in (**e**) CD45^+^CD3^-^CD11b^+^Ly6g^+^Ly6c^-^ neutrophils or (**f**) CD45^+^CD3^-^CD11b^+^Ly6g^-^Ly6c^+^ monocytes in SKP605 sg*Eed* tumors with indicated treatment. 1×10^6^ of SKP605 sg*Eed* cells were subcutaneously implanted into the right flank of WT or *Nos2^-/-^* C57BL/6J mice, IT co-injections of MQ833 and G-CSF, were given twice a week once the tumors were established. **g**, Kaplan-Meier survival curve of WT or *Nos2*^-/-^ mice bearing SKP605 sg*Eed* tumor. **h,** Tumor volumes WT or *Nos2*^-/-^ mice over days post treatment. **i,** working model of the synergistic effect of MQ833 and G-CSF.

### Co-injection of MQ833 and human G-CSF enhances the therapeutic efficacy in MPNST by promoting iNOS expression in tumor-infiltrating neutrophils

Given the importance of neutrophils in MQ833-induced antitumor effects in the MPNST tumors, we investigated whether boosting neutrophil responses by co-administration of Granulocyte Colony-Stimulating Factor (G-CSF) would enhance therapeutic efficacy. G-CSF acts primarily on the bone marrow, where it stimulates the proliferation, differentiation, and survival of precursor cells in the neutrophil lineage. It also enhances the release of mature neutrophils from the bone marrow into the bloodstream. In addition, G-CSF also activates neutrophils, enhancing their functions. G-CSF is used clinically to increase the number of neutrophils in patients with neutropenia, especially those undergoing chemotherapy or bone marrow transplantation^39^.

We first evaluated the syngeneic effect of IT MQ833 and G-CSF in PRC2-wt MPNST model, in which MQ833 monotherapy is less effective than in PRC2-loss MPNST. Briefly, SKP605 sg*Con* cells were implanted subcutaneously into the right flanks of C57/BL6 mice. When the tumors were established, they were injected with either MQ833 alone (8 x 10^7^ pfu), human G-CSF (3 µg), or a combination of MQ833 plus human G-CSF, or PBS twice weekly (Fig. 6a). Human G-CSF was used here because we previously observed that it had similar effects in boosting MQ833-induced tumor-killing as the murine version (Supplementary Fig. 5a-c). The Kaplan-Meier survival and the tumor growth curves of SKP605 sg*Con* tumor-bearing mice showed that the co-administration of MQ833 with G-CSF twice weekly resulted in a dramatic improvement in antitumor immunity compared with either agent alone in the MPNST model (Fig. 6b, c). These results demonstrated that co-administration of MQ833 and human G-CSF enhances the anti-tumor effect of MQ833 in MPNST.

We hypothesized that co-administration of MQ833 with G-CSF would increase neutrophil recruitment into the injected tumors and activate neutrophil functions. To test that, we implanted SKP605 sg*Con* cells subcutaneously into the right flanks of wild type C57BL/6J mice. After tumors were established, they were injected with MQ833 alone, human G-CSF alone, MQ833 plus human G-CSF, or PBS twice three days apart. The injected tumors were harvested two days after the second treatment. After obtaining single cell suspension, the tumor samples were stained with various markers, including CD45, CD11b, Ly6G, Ly6C, CD68, and iNOS to evaluate neutrophil, macrophage, and monocyte recruitment and activation after either MQ833 monotherapy or combination therapy with human G-CSF (Fig. 6d, e). Flow cytometry results showed that IT MQ833 induced the recruitment of neutrophils into the injected tumors and increased expression of the activation marker iNOS in neutrophils, and that the combination of IT MQ833 and human G-CSF significantly enhanced the expression of iNOS and CD11b expression in neutrophils compared with IT MQ833 alone (Fig. 6d, e). IT MQ833 and IT MQ833 plus human G-CSF resulted in similar increases in iNOS^+^ monocytes in the injected tumors (Fig. 6d, f). These results demonstrated that the co-administration of MQ833 with human G-CSF resulted in increased neutrophil recruitment and activation in the injected tumors.

### iNOS contributes to the therapeutic efficacy of MQ833 plus human G-CSF

We have previously shown that iNOS is important for MQ833-induced antitumor effects in murine B16-F10 melanoma and MC38 colon cancer models^30^. Given that the co-administration of MQ833 and human G-CSF resulted in a dramatic increase in iNOS^+^ neutrophils, we tested whether iNOS also plays a role in MQ833 plus human G-CSF-induced tumor killing in the MPNST model. We implanted SKP605 sg*Eed* cells subcutaneously into the right flanks of wild type and Nos2^-/-^ C57BL/6J mice. Once the tumors reached 4 mm in diameter, they were treated with a combination of MQ833 and human G-CSF twice weekly. The absence of *Nos2* did not affect the initial tumor sizes prior to treatment. During a 26-day treatment course, 5 out of 11 *Nos2*^-/-^mice died, while all WT mice survived (Fig. 6g, h). This result demonstrated the pivotal role of iNOS in the antitumor effects induced by MQ833 plus human G-CSF. It is likely that nitric oxide production by both activated neutrophils and macrophages/monocytes contribute to tumor killing.

## Discussion

Our previous published work has established recombinant MVA as a novel cancer immunotherapy platform^22,23,29,30^. It is based on rational design to activate the nucleic acid-sensing pathways of the host immune cells and tumor cells by deleting viral immune evasions genes and expressing transgenes that modulate immunosuppressive TME. It has several distinct features, including safety, potent induction of type I IFN, a large capacity for payloads leading to extensive remodeling of TME, stability of the viral vector, and established manufacturing processes. Our current investigation marks an extension of the therapeutic application of MQ833, transitioning from its well-established efficacy in immunogenic tumors, such as the B16-F10 melanoma and MC38 colon cancer models, to the challenging landscape of “immune-desert” PRC2-loss MPNST, known for their resistance to immune checkpoint blockade (ICB) therapy. IT delivery of MQ833 enhanced tumor immune infiltrates and sensitized PRC2-loss tumors to ICB. The synergy achieved through the co-administration of MQ833 and human G-CSF further amplifies the therapeutic potential.

Mechanistically, IT MQ833-induced antitumor immunity requires the collaborative actions of activated neutrophils, macrophages, CD4^+^, and CD8^+^ T cells (Fig. 6i). scRNA-seq analysis of MQ833-treated tumors revealed a large influx of neutrophils and their differentiation into distinct and heterogenous neutrophil populations upon IFN and other signals. Myeloid cell remodeling also involves the recruitment of monocytes into the injected tumors and differentiation into antitumor M1 macrophages, as well as the depletion of resident pro-tumor M2 macrophages. Together with the depletion of Tregs, and the expansion of ISG^+^ CD4^+^ T cells, effector CD8^+^ T cells, and proliferating CD4^+^ and CD8^+^ T cells, the “immune-desert” MPNST tumors are destroyed by MQ833 monotherapy, or they are sensitized to ICB therapy (Fig. 6i).

MQ833 was designed to induce potent induction of type I IFN in both immune and tumor cells by deleting E3L, E5R, WR199 from the MVA genome. Using *IRF3*^-/-^*IRF7*^-/-^ and *Stat2*^-/-^ mice, our results showed that type I IFN production and IFNAR signaling is critical for MQ833-induced antitumor effects in MPNST tumors. Using IFNAR1 conditional knockout mice in CD4^+^ and CD8^+^ T cells, our results demonstrate that IFNAR1 signaling on CD4^+^ and CD8^+^ T cells are crucial for their expansion and activation. This correlates with the expansion of ISG^+^ CD4^+^ T cells and effector CD8^+^ T cells revealed by scRNA-seq analysis of CD45^+^ immune cells in MPNST tumors treated with MQ833. CD4^+^ T cells were found to be crucial for MQ833-induced antitumor effects in the MPNST tumors but were dispensable for MQ833-induced tumor-killing in the B16-F10 melanoma model. We speculate that upon IFN stimulation, ISG^+^CD4^+^ T cells undergo expansion and activation, leading to tumor indirect killing through MHC-II-dependent and MHC-II-independent mechanisms^40–43^.

We have previously reported the role of type I IFN on the recruitment and polarization of monocytes upon MQ833 treatment in the B16-F10 melanoma model^30^. Here, using a neutrophil-specific conditional knockout of IFNAR1, we showed that IFNAR signaling on neutrophils are critical for neutrophil-mediated tumor killing. This aligns with our scRNA-seq analysis that identified a subset of neutrophils expressing interferon-stimulated genes (ISGs). Interestingly, these ISG-expressing neutrophils do not coincide with those expressing nitric oxide synthase 2 (Nos2), suggesting that ISG^+^ neutrophils may also execute anti-tumor functions independently of iNOS. It has been recently reported that neutrophils expressing ISG signature mediate effective immunotherapy, including anti-CD40 agonist antibody^44^.

PRC2-loss MPNST exhibit an ‘immune-desert’ profile and show resistance to ICB therapy. Contrary to expectations, our data reveals that PRC2-deficient tumors (SKP605 sg*Eed*) respond more favorably to both monotherapy with MQ833 and its combination with ICB, compared to PRC2-wt (wild-type) tumors (SKP605 sg*Con*). This heightened sensitivity to MQ833 is associated with increased interferon-beta (IFN-β) production by the PRC2-deficient tumor cells. Single-cell RNA sequencing (scRNA-seq) of CD45-positive immune cells within SKP605 tumors indicates a reduced presence of monocytes/M1 macrophages, an increased prevalence of regulatory T cells (Tregs), and a higher number of exhausted Pdcd1^+^Lag3^+^ CD8^+^ T cells in PRC2-loss tumors, aligning with their non-immunogenic nature. IT MQ833 modifies the suppressive TME to a more immune-activating one, thus synergizing with ICB therapy. Given that over 80% of high-grade MPNSTs exhibit PRC2 loss, our findings provide a compelling scientific basis for initiating clinical trials to evaluate the efficacy of MQ833, alone or in conjunction with ICBs, in treating this aggressive form of cancer.

The combination of MQ833 with human G-CSF has generated superior antitumor effects compared to either agent alone. Human G-CSF promotes neutrophil recruitment and iNOS expression in neutrophils. Moreover, iNOS plays a crucial role in the antitumor effects induced by the combination of MQ833 and human G-CSF. These results provide further evidence that neutrophils can be mobilized for antitumor activity. Such findings are anticipated to significantly influence the design of clinical trials for treating MPNST tumors in patients. Human G-CSF is commonly prescribed for patients with neutropenia. Future studies will assess whether the systemic delivery of G-CSF could also have synergistic effects when used with MQ833 delivered intratumorally.

In summary, our innovative design of a recombinant non-replicative Modified Vaccinia Ankara (MVA) virus for cancer immunotherapy has opened new avenues to explore how the immunosuppressive TME can be altered. We have determined that the induction of type I IFN by the immune-stimulating recombinant MVA within the TME, along with IFNAR signaling, is crucial for the activation of both innate and adaptive immunity, which leads to tumor destruction. Our research indicates that cytotoxic CD4+ T cells significantly contribute to tumor clearance in the malignant peripheral nerve sheath tumor (MPNST) model, but this is not the case in the melanoma model. Furthermore, we have discovered the importance of IFNAR signaling in neutrophils for their anti-tumor activity, and that human granulocyte colony-stimulating factor (G-CSF) can amplify their tumor-destroying properties. Lastly, our first-generation recombinant MVA (MQ710) has progressed to a Phase I clinical trial for solid tumors (NCT05859074), promising to yield critical insights into the human response to virus-based cancer immunotherapy and inform future therapeutic designs.

## Methods

### Mice

Female C57BL/6J mice between 6 and 8 weeks of age were purchased from the Jackson Laboratory (Strain #000664) were used for in vivo experiments. These mice were maintained in the animal facility at the Sloan Kettering Institute. All procedures were performed in strict accordance with the recommendations in the Guide for the Care and Use of Laboratory Animals of the National Institutes of Health. The protocol was approved by the Committee on the Ethics of Animal Experiments of Sloan Kettering Cancer Institute. *Rag1^-/-^* (Strain #002216), *Stat2^-/-^*(Strain #023309), *Mrp8*-cre^45^ (Strain #021614) and *Nos2^-/-^*(Strain #002609) mice were purchased from Jackson Laboratory. *Irf3^-/-^ Irf7^-/-^, Mrp8^cre^Ifnar^fl/fl^, Cd4^cre^Ifnar1^fl/fl^* were bred in our lab.

### Cell lines

BHK-21 (baby hamster kidney cell, ATCC CCL-10) cells were cultured in Eagle’s minimal essential medium containing 10% fetal bovine serum (FBS), 0.1 mM nonessential amino acids, penicillin, and streptomycin. The murine MPNST cell line, SKP605 (*Nf1^−/−^, Cdkn2a/b^−/−^*), was generated in Ping Chi’s laboratory (Memorial Sloan Kettering Cancer Center) and they were cultured in Dulbecco’s Modified Eagle Medium (DMEM) supplemented with l-glutamine (2 mM), penicillin (100 U/mL), streptomycin (100 μg/mL), and 10% heat-inactivated FBS in 5% CO2 at 37°C. The murine melanoma cell line B16-F10 was originally obtained from I. Fidler (MD Anderson Cancer Center). The B16-F10 cell line lacking B2m gene were generated by using CRISPR-cas9 technology as described^30^. The B16-F10 cell line was maintained in RPMI-1640 medium supplemented with 10% FBS, 0.05 mM 2-mercaptoethanol, penicillin, and streptomycin in 5% CO2 at 37°C.

### Viruses

The MVA virus was a kind gift from Gerd Sutter (University of Munich). MQ833 was generated as described^30^. Viruses were propagated in BHK-21 cells and purified through a 36% sucrose cushion.

### ELISA

For murine IL-12-p70 and IFN-β, BHK-21, SKP605 sg*Con* and sg*Eed* 16 cells were infected with MQ833 at a MOI of 10 for 1 hour. Mock-infected cells were used as a control. The inoculum was removed, and the cells were washed with PBS twice and incubated with fresh medium for 16 hours. Supernatant was collected from the culture. IL-12 and IFN-β levels were determined using Duoset ELISA kit (R&D) following manufacturer’s instruction.

### Western blot analyses

BHK-21, SKP605 sg*Con* and sg*Eed* were infected with MQ833 at a MOI of 10 for 1 hour. Mock-infected cells were used as a control. Cells were then washed with PBS twice and cultured with fresh medium. Cells were lysed in RIPA lysis buffer supplemented with 1× Halt™ Protease and Phosphatase Inhibitor Cocktail at indicated time points. Protein samples were separated by SDS-PAGE and then transferred to nitrocellulose membrane. Protein levels of cGAS, IRF-3, p-IRF-3, STING, p-STING and H3K27me3 were measured, and GAPDH was used as an internal control. Primary antibodies specific for cGAS (1:1000, CST, 31659), IRF3 (1:1000, CST, 4302), phospho-IRF3 (1:1000, CST, 4947) and H3K27me3 (1:1000, CST, 9733). Anti-mouse HRP-linked IgG antibody was used as a secondary antibody (1:5000, CST, 7074 or 7076) were used. GAPDH antibody (1:5000, CST, 2118) were used as loading controls. Detection was performed using SuperSignal^TM^ Substrates (Thermo Fisher, 34577 or 34095).

### Tumor processing and flow cytometry staining

Tumor samples were collected in cold RPMI and thoroughly chopped into small pieces with surgical scissor. Then we added Liberase TL (1.67 Wünsch U/ml) and DNase I (0.2 mg/ml) to the tube and incubated in 37 °C shaker for 20-30 minutes. Digested tumors were then transferred to GentleMACs C Tube and homogenized using the Gentle MACs Octo Dissociator (Miltenyi Biotec) according to manufacturer’s instruction. Samples were then filtered through 70 um filter and quenched with 10 ml cold PBS and spun down. Cells were then washed with MACS buffer (Miltenyi Biotec) twice, and stained with Fc block (BD), viability dye eFluor506 (eBioscience), and cell surface antibodies diluted in MACS buffer for 30 minutes on ice in the dark, and subsequently fixed and permeabilized using the Foxp3 fixation and permeabilization kit (Thermo Fisher). Cells were then incubated with intracellular antibodies diluted in permeabilization buffer for 30 minutes or overnight.

The following antibodies were used for flow cytometric staining: CD45 Pacific blue (clone 30-F11, Biolegend), CD3 BUV395 (clone 145-2C11, BD), CD4 BUV737 (clone RM4-5, BD), CD8 Alexa Fluor 700 (clone 53-6.7, Biolegend), OX40 BV605 (clone OX-86, Biolegend), TIM3 BV711 (clone RMT3-23, Biolegend), PD-1 AF647 (clone RMPI-30, BD), KLRG1 APC-Cy7 (clone 2F1/KLRG1, Biolegend), Foxp3 PerCP-Cy5.5 (clone FJK-16s, Invitrogen), Ki67 BV786 (clone B56, BD), TOX PE (clone REA473, Miltenyi Biotec), Granzyme B PE-Texas red (clone GB11, Invitrogen), CD11b APC-eFluor780 (clone M1/70, eBioscience), Ly6G PE-Cy7 (clone 1A8, BD), F4/80 APC (clone BM8, Biolegend), CD68 BV421 (clone FA-11, Biolegend), iNOS PE (clone W16030C, Biolegend). After staining, cells were washed and resuspended with MACS buffer and transferred into FACS tubes with 70 um filter caps. Single cell suspensions were run on Cytek Aurora analyzer, and the data was analyzed with Flowjo.

### Tumor challenge and treatment

For survival experiments, flow cytometric analysis and single cell RNA-seq, 1 ×10^6^ SKP sg*Con* or sg*Eed* cells were implanted subcutaneously into the shaved skin on the right flanks of WT C57BL/6J mice. Once the tumors are 3 mm in diameter or larger, they were injected with 8 ×10^7^ PFU of MQ833. Viruses were injected twice weekly as specified in each experiment and tumor sizes were measured twice a week. For combination therapy with ICBs, the following antibodies were injected interperitoneally twice weekly: anti-CTLA-4 (100 µg per mouse) and anti-PD-1 (200 µg per mouse). Tumor volumes were calculated according to the following formula: l (length) × w (width) × h (height)/2. Mice were euthanized for signs of distress or when the diameter of the tumor reached 10 mm.

### Cell sorting for tumor infiltrating immune cells

Tumors were harvested and single cell suspensions were prepared. Cells were then stained with Cell Viability Dye (eFlour506, brand) and anti-CD45 antibody (brand). CD45^+^ live cells were then sorted using B&D Aria cell sorter.

### Single-cell RNA-sequencing and data analysis

#### Library preparation and sequencing

CD45^+^ populations from tumors were purified by FACS sorting. We performed single-cell 5′ gene expression profiling on the single-cell suspension using the Chromium Single Cell V(D)J Solution from 10x Genomics according to the manufacturer’s instructions. Cell-barcoded 5′ gene expression libraries were sequenced on an Illumina NovaSeq6000 sequencer with pair-end reads.

#### Gene expression UMI counts matrix generation

The sequencing data were primarily analyzed by the 10× cellranger pipeline (v3.0.2) in two steps. In the first step, cellranger mkfastq demultiplexed samples and generated fastq files; and in the second step, cellranger count aligned fastq files to the reference genome and extracted gene expression UMI counts matrix. In order to measure both human and viral gene expression, we built a custom reference genome by integrating the MVA virus genome into the 10× pre-built mouse reference using cellranger mkref. The MVA virus genome was downloaded from NCBI.

#### Single-cell RNA-seq data analysis

All cells expressing <200 or >6,000 genes were removed as well as cells that contained >5% mitochondrial counts. Samples were merged and normalized. The default parameters of Seurat were used, unless mentioned otherwise. Briefly, 2,000 variable genes were identified for the clustering of all cell types and principal component analysis (PCA) was applied to the dataset to reduce dimensionality after regressing for the number of UMIs (counts). The top 12-15 most informative principal components (PCs) were used for clustering and Uniform Manifold Approximation and Projection for dimension reduction (UMAP). To characterize each cluster, we applied both the *FindAllMarkers* and *FindMarkers* procedure in Seurat, which identified markers using log fold changes (FC) of mean expression. To identify differentially expressed genes between two groups of clusters, *FindMarkers* functions in Seurat and Enhanced Volcano R package (v1.8.0) were used. For subsequent T cell analysis, we first extracted all the CD3^+^ cells from the original dataset and repeated the clustering procedures in the T cell subset.

### Purification of anti-CSF-1 monoclonal antibody

The anti-CSF-1 monoclonal antibody (clone 5A1) was purified from cell supernatant of hybridoma (ATCC, #CRL-2702) in Dr. Ming Li’s laboratory (Memorial Sloan Kettering Cancer Center). Briefly, hybridoma cells were cultured in Hybridoma-SFM medium (Thermo #12045076) for 3 weeks to produce antibody. Cell supernatant was collected to do antibody purification by using LigaTrap Rat IgG purification column (LigaTrap Technologies #LT-138) according to the manufacturer’s procedure. Antibody quality and concentration were examined by SDS-PAGE gel electrophoresis and nanodrop, respectively.

### In vivo antibody depletion experiment

Immune cell subsets were depleted by administering 200 μg monoclonal antibodies i.p. twice weekly starting 1 day prior to the first viral injection as indicated, continued until the animals either died, were euthanized, or were completely clear of tumors: anti-CD8-α for CD8^+^ T cells (clone 2.43, BioXCell), anti-CD4 for CD4^+^ T cells (clone GK1.5, BioXCell), anti-NK1.1 for NK cells (clone PK136, BioXCell). 500 μg of anti-Ly6G (clone 1A8, BioXCell) were given for neutrophil depletion. 500 μg of anti-CSF1 (provided by Ming Li’s laboratory) were given every 5 days for macrophage depletion. Depletion efficacies of CD8 T cells, CD4 T cells, NK cells and neutrophils were confirmed by flow cytometry of peripheral blood.

### Statistical analysis

Two-tailed unpaired Student’s t test was used for comparisons of two groups in the studies. Survival data were analyzed by log-rank (Mantel-Cox) test. The p values deemed significant are indicated in the figures as follows: *, p < 0.05; **, p < 0.01; ***, p < 0.001; ****, p < 0.0001. The numbers of animals included in the study are discussed in each figure legend.

### Lead Contact

Further information and requests for resources and reagents used in this study should be directed to and will be fulfilled by the lead contact and the corresponding author, Liang Deng.

### Materials availability

Materials generated in our laboratory are available upon request.

### Data and code availability

The scRNA-seq data reported in this study will be deposited in the Gene Expression Omnibus database (GEO) and will make it available to the public after the manuscript is accepted for a publication after peer-review.

## Acknowledgements

We thank the Flow Cytometry Core Facility at the Sloan Kettering Institute and Genomic Resources Core Facility Weill Cornell Medical College. This work was supported by the Society of Memorial Sloan Kettering cancer center (MSKCC) research grant (L.D.), MSKCC Technology Development Fund (L.D.), Parker Institute for Cancer Immunotherapy at MSKCC (L.D.), Geoffrey Beene Grant (P.C., L.D., M.O.L), Cycle for Survival (L.D., P.C., M.O.L.), Sponsored Research Award from IMVAQ Therapeutics (L.D.), 1R01CA280657 (P.C., L.D.), and Sarcoma Grant (L.D., P.C., M.O.L.). Our working model figure was created with BioRender.com.

## Author Contributions

Author contributions: L.D., Y.Q.W., and S.L. were involved in all aspects of this study, including designing and performing experiments, data analysis and interpretation, and manuscript writing. L.D., J.Y., and P.C. established the viral-treatment model in MPNST tumors. S.L. and Y.Q.W. performed scRNA-seq analysis. S.B.T. generated the T cell-specific IFNAR1 conditional knockout mice and *Irf3^-/-^Irf7^-/-^*. S.L and Y.Q.W. generated the neutrophil specific IFNAR1 conditional knockout mice. S.B.T. and B.S. assisted with some experiments. L.L.J. produced the anti-CSF1 antibody. M.O.L. and P.C. assisted in some experimental design, data interpretation, and manuscript preparation. L.D. provided overall supervision of the study.

