## Supplementary Materials for "Activating neutrophils by co-administration of immunogenic recombinant modified vaccinia virus Ankara and granulocyte colony-stimulating factor for the treatment of malignant peripheral nerve sheath tumor"

**This file contains:**

- Supplementary Fig. 1
- Supplementary Fig. 2
- Supplementary Fig. 3
- Supplementary Fig. 4
- Supplementary Fig. 5

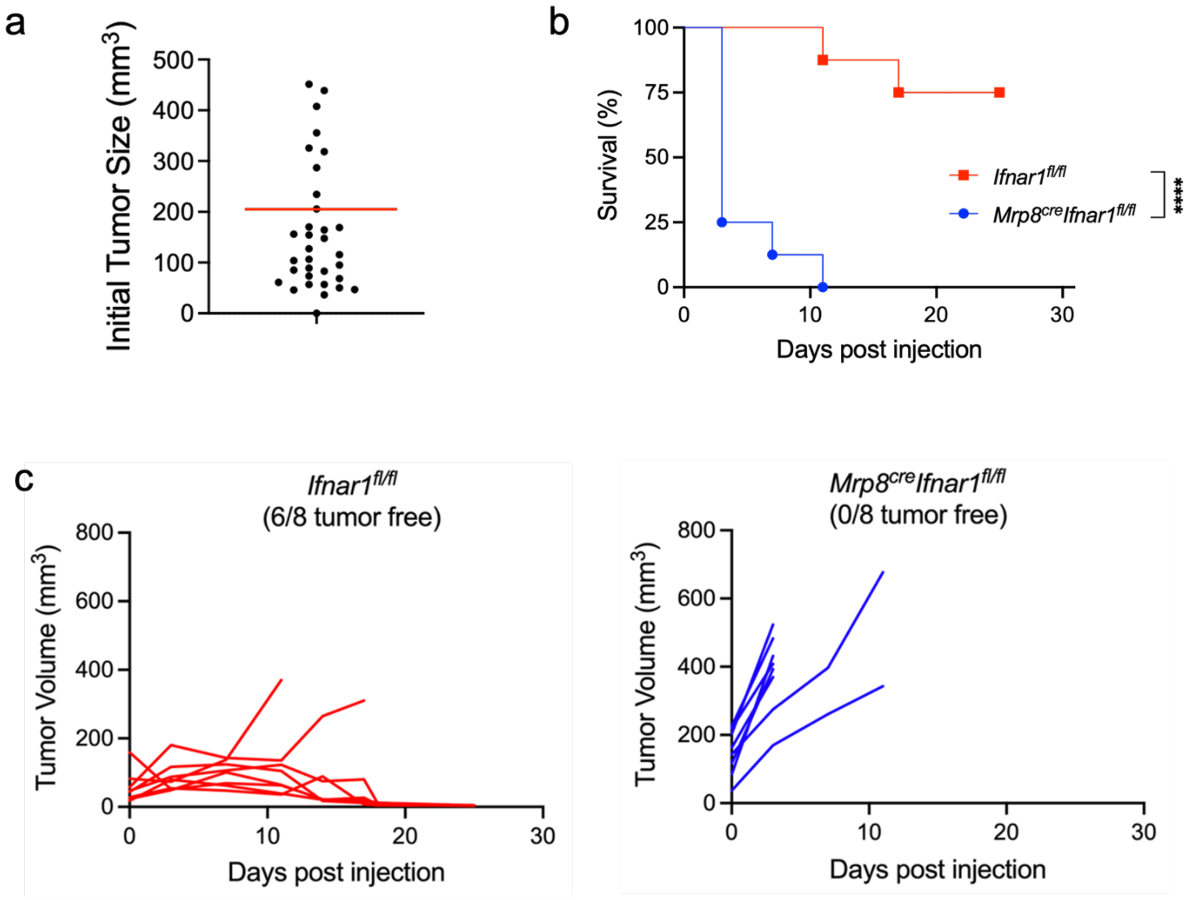
a
Initial Tumor Size (mm3)
500
400
300
200
100
0
b
Survival (%)
100
75
50
25
0
Ifnar1fl/fl
Mrp8creIfnar1fl/fl
****
0
10
20
30
Days post injection
c
Ifnar1fl/fl
(6/8 tumor free)
Tumor Volume (mm3)
800
600
400
200
0
Days post injection
Mrp8creIfnar1fl/fl
(0/8 tumor free)
Tumor Volume (mm3)
Days post injection

**Supplementary Fig.1| IFNAR signaling in neutrophils plays a role in IT MQ833-induced antitumor effects**. $5x10^5$ B16-F10 cells were intradermally implanted into the right flank of *Mrp8*$^{cre}$*Ifnar*$^{fl/fl}$ or *Ifnar*$^{fl/fl}$ C57BL/6J mice. Intratumoral injection of MQ833 were given twice a week once the tumors were established. Tumor size and mice survival were monitored. **a,** Initial tumor volume for each group. **b,** Kaplan-Meier survival curve of B16-F10 tumor bearing mice treated with MQ833 virus. PBS was used as a control. Survival data were analyzed by log-rank (Mantel-Cox) test (n=8, ****$P < 0.0001$). **c,** Tumor volumes of mice injected with MQ833 or PBS over days post treatment.

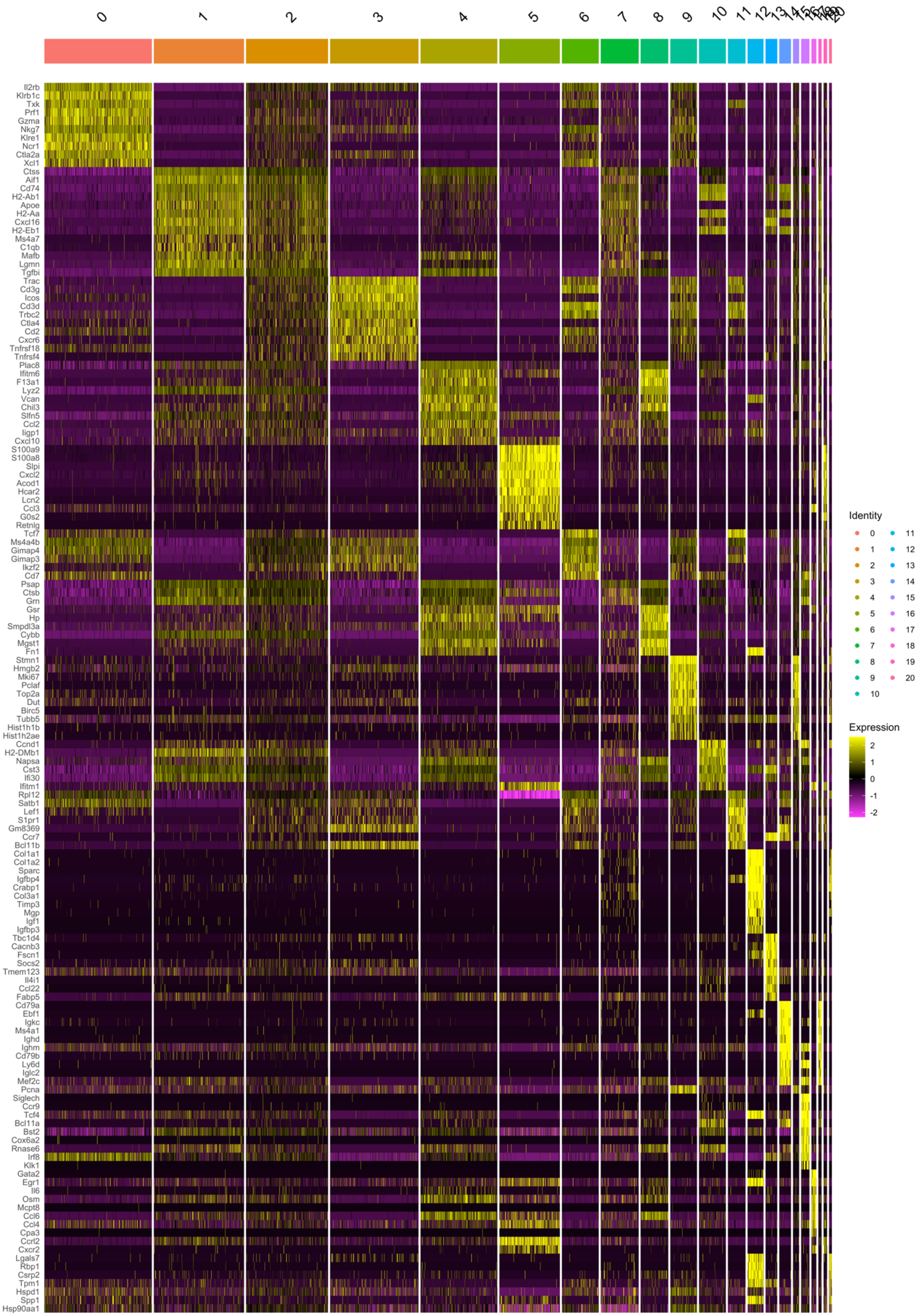
0
1
2
3
4
5
6
7
8
9
10
11
12
Il2rb
Klrb1c
Txk
Prf1
Gzma
Nkg7
Klre1
Ncr1
Ctla2a
Xcl1
Ctss
Aif1
Cd74
H2-Ab1
Apoe
H2-Aa
Cxcl16
H2-Eb1
Ms4a7
C1qb
Mafb
Lgmn
Tgfbi
Trac
Cd3g
Icos
Cd3d
Trbc2
Ctla4
Cd2
Cxcr6
Tnfrsf18
Tnfrsf4
Plac8
Ifitm6
F13a1
Lyz2
Vcan
Chil3
Slfn5
Ccl2
Iigp1
Cxcl10
S100a9
S100a8
Slpi
Cxcl2
Acod1
Hcar2
Lcn2
Ccl3
G0s2
Retnlg
Tcf7
Ms4a4b
Gimap4
Gimap3
Ikzf2
Cd7
Psap
Ctsb
Grn
Gsr
Hp
Smpdl3a
Cybb
Mgst1
Fn1
Stmn1
Hmgb2
Mki67
Pclaf
Top2a
Dut
Birc5
Tubb5
Hist1h1b
Hist1h2ae
Ccnd1
H2-DMb1
Napsa
Cst3
Ifi30
Ifitm1
Rpl12
Satb1
Lef1
S1pr1
Gm8369
Ccr7
Bcl11b
Col1a1
Col1a2
Sparc
Igfbp4
Crabp1
Col3a1
Timp3
Mgp
Igf1
Igfbp3
Tbc1d4
Cacnb3
Fscn1
Socs2
Tmem123
Il4i1
Ccl22
Fabp5
Cd79a
Ebf1
Igkc
Ms4a1
Ighd
Ighm
Cd79b
Ly6d
Iglc2
Mef2c
Pcna
Siglech
Ccr9
Tcf4
Bcl11a
Bst2
Cox6a2
Rnase6
Irf8
Klk1
Gata2
Egr1
Il6
Osm
Mcpt8
Ccl6
Ccl4
Cpa3
Ccrl2
Cxcr2
Lgals7
Rbp1
Csrp2
Tpm1
Hspd1
Spp1
Hsp90aa1
Identity
Expression
2
1
0
-1
-2

**Supplementary Fig. 2| Heatmap of top 10 genes in CD45$^+$ cells in SKP605 sg*Con* and sg*Eed* tumor treated with MQ833 or PBS.**

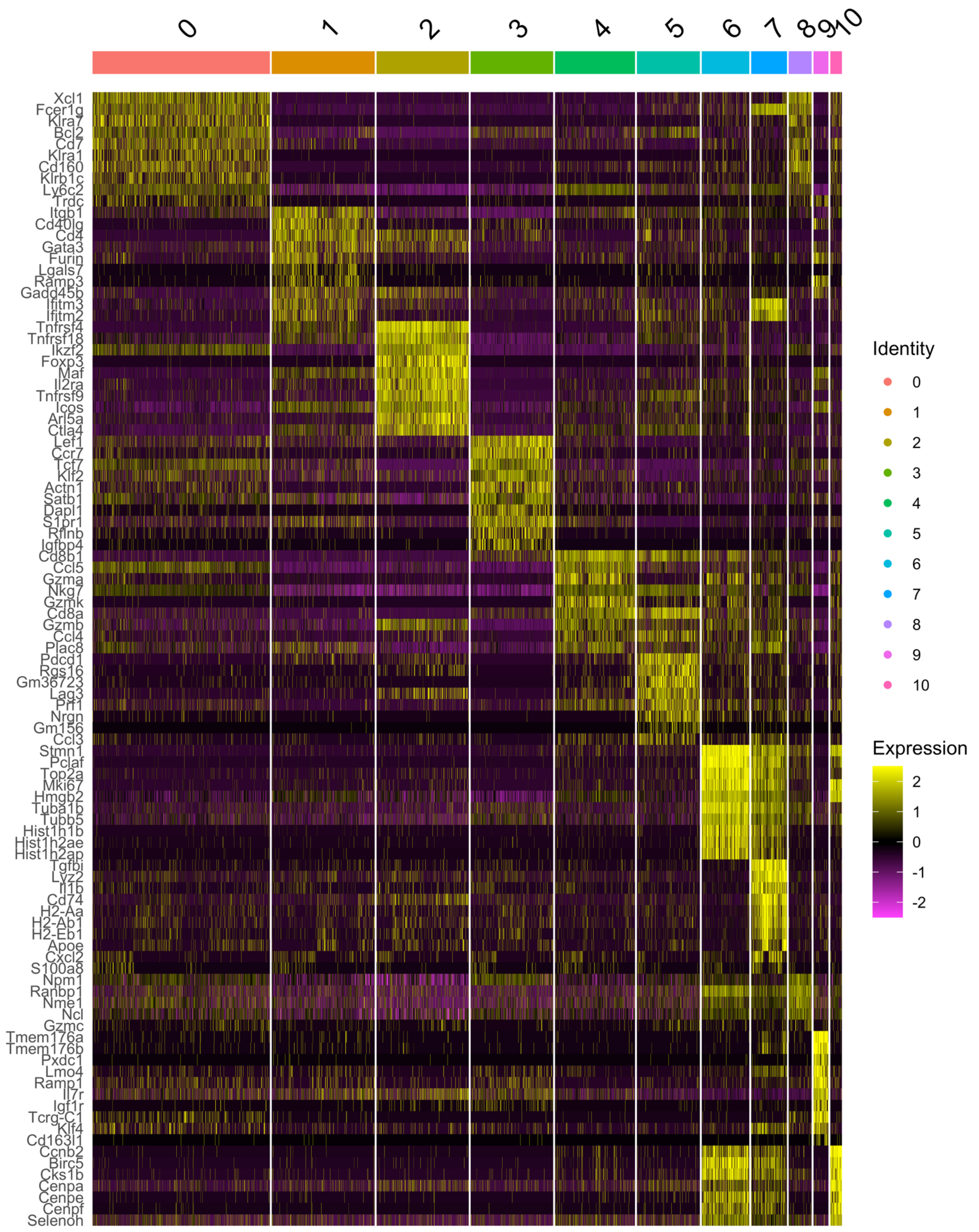
0
1
2
3
4
5
6
7
8
9
10
Xcl1
Fcer1g
Klra7
Bcl2
Cd7
Klra1
Cd160
Klrb1c
Ly6c2
Trdc
Itgb1
Cd40lg
Cd4
Gata3
Furin
Lgals7
Ramp3
Gadd45b
Ifitm3
Ifitm2
Tnfrsf4
Tnfrsf18
Ikzf2
Foxp3
Maf
Il2ra
Tnfrsf9
Icos
Arl5a
Ctla4
Lef1
Ccr7
Tcf7
Klf2
Actn1
Satb1
Dapl1
S1pr1
Rflnb
Igfbp4
Cd8b1
Ccl5
Gzma
Nkg7
Gzmk
Cd8a
Gzmb
Ccl4
Plac8
Pdcd1
Rgs16
Gm36723
Lag3
Prf1
Nrgn
Gm156
Ccl3
Stmn1
Pclaf
Top2a
Mki67
Hmgb2
Tuba1b
Tubb5
Hist1h1b
Hist1h2ae
Hist1h2ap
Tgfbi
Lyz2
Il1b
Cd74
H2-Aa
H2-Ab1
H2-Eb1
Apoe
Cxcl2
S100a8
Npm1
Ranbp1
Nme1
Ncl
Gzmc
Tmem176a
Tmem176b
Pxdc1
Lmo4
Ramp1
Il7r
Igf1r
Tcrg-C1
Klf4
Cd163l1
Ccnb2
Birc5
Cks1b
Cenpa
Cenpe
Cenpf
Selenoh
Identity
0
1
2
3
4
5
6
7
8
9
10
Expression
2
1
0
-1
-2

**Supplementary Fig. 3| Heatmap of top 10 genes of T cell subclusters in SKP605 sg*Con* and sg*Eed* tumors treated with MQ833 or PBS.**

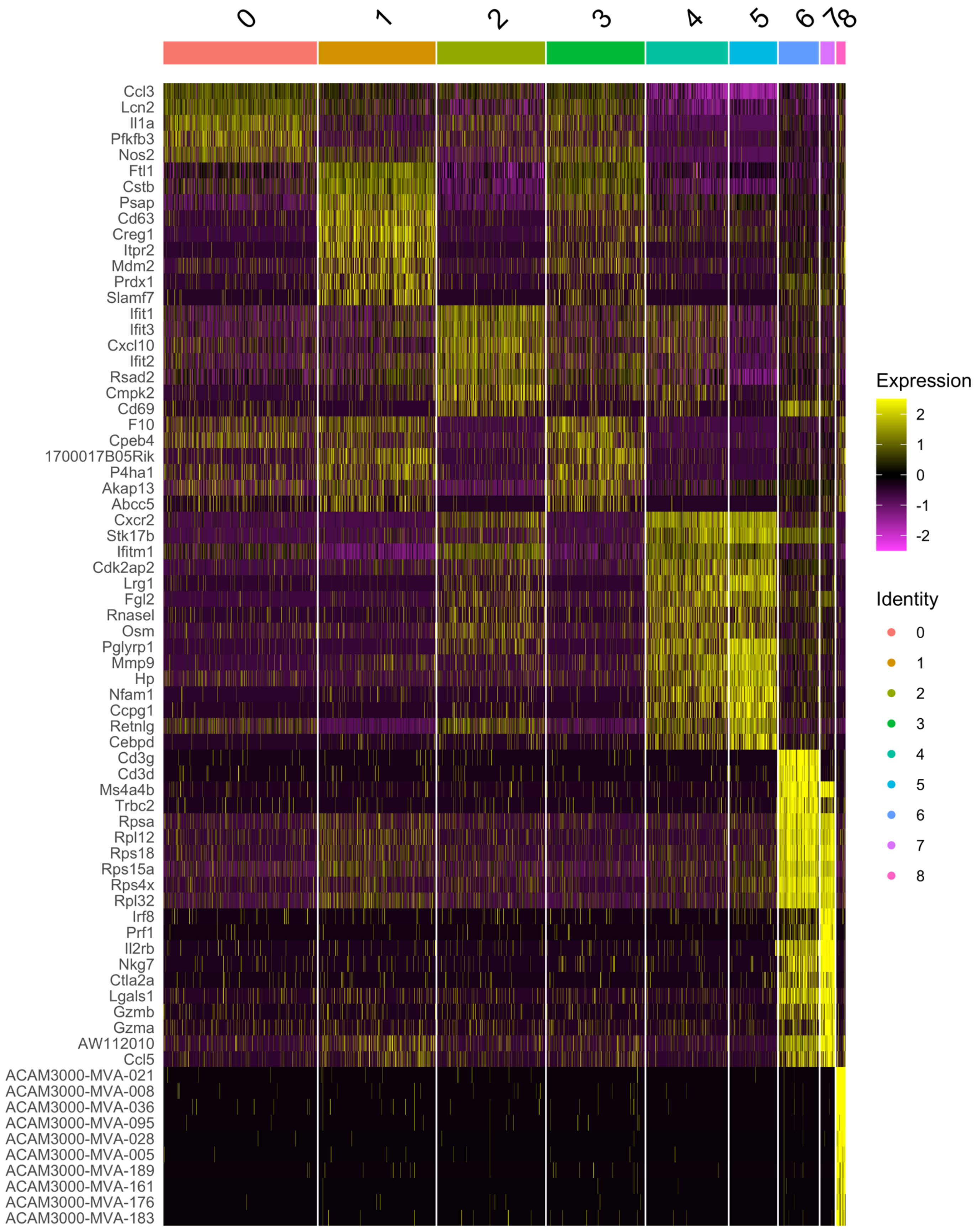
0
1
2
3
4
5
6
7
8
Ccl3
Lcn2
Il1a
Pfkfb3
Nos2
Ftl1
Cstb
Psap
Cd63
Creg1
Itpr2
Mdm2
Prdx1
Slamf7
Ifit1
Ifit3
Cxcl10
Ifit2
Rsad2
Cmpk2
Cd69
F10
Cpeb4
1700017B05Rik
P4ha1
Akap13
Abcc5
Cxcr2
Stk17b
Ifitm1
Cdk2ap2
Lrg1
Fgl2
Rnasel
Osm
Pglyrp1
Mmp9
Hp
Nfam1
Ccpg1
Retnlg
Cebpd
Cd3g
Cd3d
Ms4a4b
Trbc2
Rpsa
Rpl12
Rps18
Rps15a
Rps4x
Rpl32
Irf8
Prf1
Il2rb
Nkg7
Ctla2a
Lgals1
Gzmb
Gzma
AW112010
Ccl5
ACAM3000-MVA-021
ACAM3000-MVA-008
ACAM3000-MVA-036
ACAM3000-MVA-095
ACAM3000-MVA-028
ACAM3000-MVA-005
ACAM3000-MVA-189
ACAM3000-MVA-161
ACAM3000-MVA-176
ACAM3000-MVA-183
Expression
2
1
0
-1
-2
Identity
0
1
2
3
4
5
6
7
8

**Supplementary Fig. 4| Heatmap of top 10 genes of neutrophil subclusters in SKP605 sg*Con* and sg*Eed* tumors treated with MQ833 or PBS.**

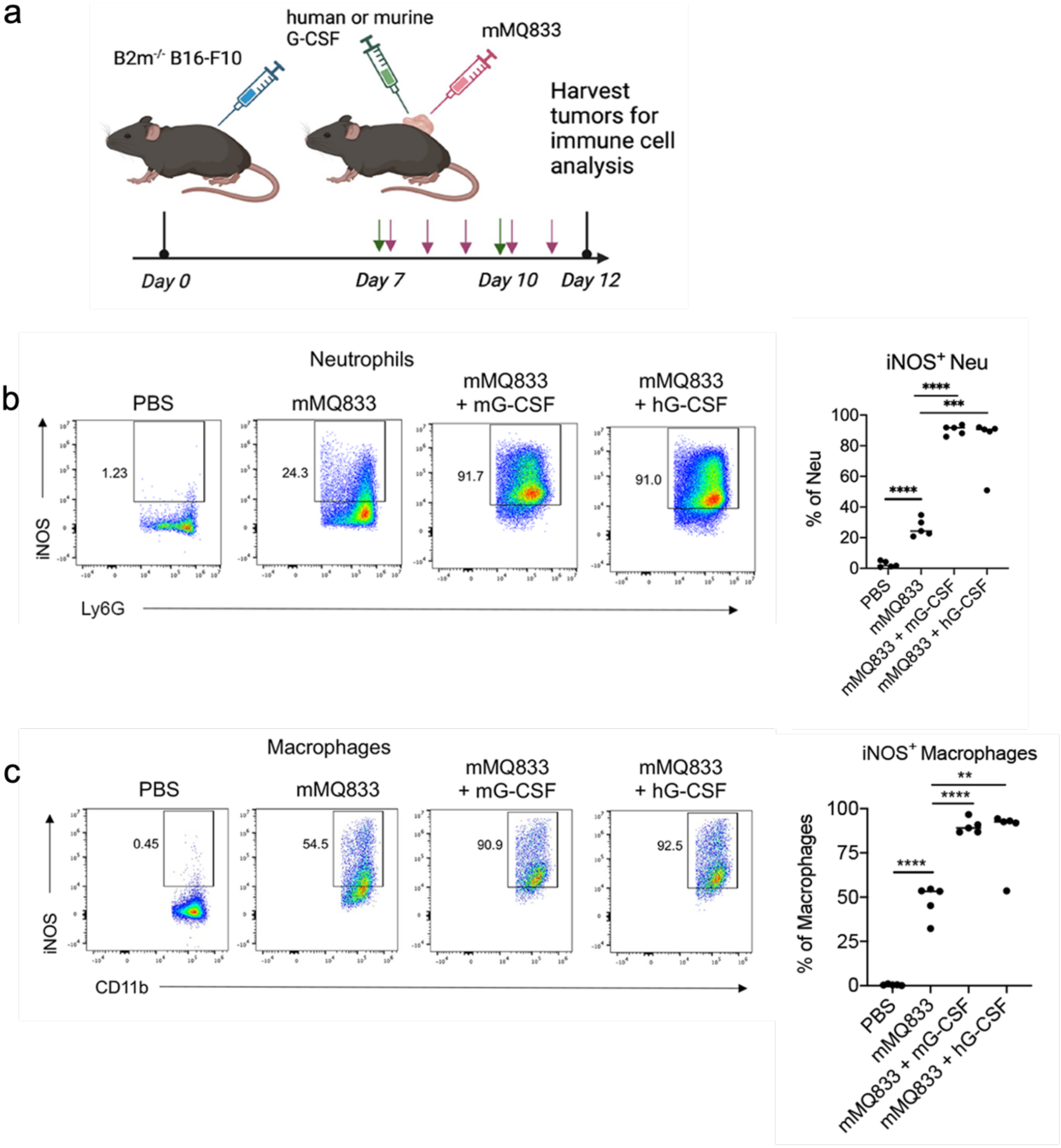
a
B2m-/- B16-F10
human or murine G-CSF
mMQ833
Harvest tumors for immune cell analysis
Day 0
Day 7
Day 10
Day 12
b
Neutrophils
PBS
mMQ833
mMQ833 + mG-CSF
mMQ833 + hG-CSF
1.23
24.3
91.7
91.0
iNOS
Ly6G
iNOS+ Neu
% of Neu
c
Macrophages
0.45
54.5
90.9
92.5
CD11b
iNOS+ Macrophages
% of Macrophages

**Supplementary Fig. 5| MQ833 combination treatment with human or murine G-CSF improves the tumor microenvironment (TME) in an Immune Checkpoint Blockade (ICB)-resistant model. a,** Schematic diagram of the experimental schedule. B2m KO B16-F10 melanoma ($6x10^5$) cells were implanted intradermally to the right flank of wild type C57BL/6J mice. After tumors were established, they were injected with murine mMQ833 ($8x10^7$ pfu) and murine granulocyte colony stimulating factor (mG-CSF) (3 µg) or human granulocyte colony stimulating factor (hG-CSF) (3 µg). mMMQ833 was injected twice, three days apart. mG-CSF or hG-CSF were given intratumorally daily for 5 consecutive days. The injected tumors were harvested two days after the second dose of mMQ833. Contour plots and percentages of $CD45^+CD3^-CD11b^+Ly6g^+$ neutrophils **(b)** and macrophages **(c)** among $CD45^+$ cells.
